# Additive Effects Dominate Legume Responses to Combined Heat and Drought Stress: A Quantitative Review

**DOI:** 10.64898/2026.08.12.744551

**Authors:** Lotus Meijer, Karine Chenu, Millicent R. Smith, Shanice Van Haeften, Victor Sadras

**Author notes:** Corresponding authors: Shanice Van Haeften, (editorial communication), Victor Sadras.

## Abstract

Concurrent exposure to heat and drought stress compromises legume productivity, yet their combined effects are rarely quantified systematically. We compiled a database of 18 studies covering seven legume species. From these, we extracted 929 physiological, biochemical, and yield-related traits and calculated actual-to-additive ratios to classify heat–drought interactions as antagonistic (ratio < 1), additive (ratio = 1), or synergistic (ratio > 1). Additive heat-drought relationships accounted for 59 % of all classifiable observations, 37% relationships were antagonistic, and 4% synergistic. The relationship varied with species, genotype, trait, and experimental conditions highlighting the complexity of combined abiotic stress effects. The results challenge the common assumption that concurrent stresses invariably exacerbate damage and underscore the need for more realistic, quantitatively defined stress treatments as well as frameworks that integrate trait-level responses into predictive models of crop growth and development. Our synthesis provides a quantitative foundation to understand legume phenotypes under the increasingly frequent co-occurrence of heat and drought stress and identifies research areas where further work is needed to improve insight into combined stress responses.

**Highlights:**

- Combined heat and drought responses were mainly additive or antagonistic.
- Evidence is biased toward few legumes and controlled environments.
- Field-based, multi-species studies are needed to identify adaptive traits.

## 1. Introduction

The frequency and intensity of heat and drought events are increasing across most agricultural regions, driven by accelerating climate variability (Ababaei & Chenu, 2020; Lake et al., 2025). These stresses are among the leading causes of yield loss in major crops and are projected to intensify further in both severity and co-occurrence under future climate scenarios (Lobell et al., 2015; Zandalinas et al., 2018; S. Zhou et al., 2024). With climate variability placing greater pressure on cropping systems, it is vital to consider the crops that support both food security and ecological sustainability. As major sources of plant-based protein, essential amino acids, and micronutrients, grain legumes are valuable to both human and animal diets (Goldstein & Reifen, 2022; Margier et al., 2018). Beyond their nutritional value, legumes contribute to sustainable agriculture through their capacity to fix atmospheric nitrogen, improve microbial diversity, and enhance soil fertility, reducing dependence on synthetic fertilisers (Yan et al., 2023; Yang et al., 2024). Their inclusion in crop rotations increases system productivity by improving the yield of both the legume and subsequent non-legume crops (Yang et al., 2024; Zhao et al., 2022). Understanding how these crops respond to simultaneous heat and drought stress is therefore useful for developing adaptive strategies that sustain productivity in warming, drying regions.

Drought and heat stress trigger distinct physiological disruptions whose individual and combined severity depend on timing, intensity, and duration of stress events (Gawinowski et al., 2025). Drought primarily restricts tissue expansion, can reduce grain set if it occurs during flowering, and limits grain size when experienced later in the season (Qiao et al., 2024; Seleiman et al., 2021). Heat stress further compromises reproduction by causing flower abortion, reducing pollen viability, and accelerating phenological development, which shortens flowering, podding and seed-filling periods (Devi et al., 2023; Djanaguiraman et al., 2013; Kumar et al., 2013; Tang et al., 2023; Van Haeften et al., 2023). Together, these effects reduce grain number and final yield (Daryanto et al., 2017; Varshney et al., 2019), while also diminishing grain quality (Farooq et al., 2017). Studies focusing on heat and drought stress have provided insights into legume adaptation but these responses are often examined in isolation (e.g. Kumar et al., 2013; Poudel, Vennam, et al., 2023). While the majority of experimental work continues to assess heat or drought independently, these stressors frequently coincide in the field (Choukri et al., 2022; Devi et al., 2023; Lobell et al., 2015; Pappula-Reddy et al., 2024; Poudel, Adhikari, et al., 2023; Vaseva et al., 2011; Z. Zhou et al., 2024). For faba bean and chickpea in Australia, new environmental characterisations have been advanced that capture the co-occurrence of multiple stresses (Lake et al., 2025; Manson et al., 2025). Combined stress can be additive (matching the sum of individual effects), antagonistic (less severe than expected), or synergistic (more severe than expected). Synergies reflect the way stresses might amplify one another: water limitation can reduce evaporative cooling and increase heat injury, while elevated temperature can exacerbate water loss and accelerate phenological development, compounding drought effects (Moore et al., 2021; Seleiman et al., 2021). Antagonistic relationships can arise from cross-talk, for example when drought triggers ABA-mediated stomatal closure but heat stimulates increased transpiration, producing opposing signals that result in a non-additive combined response dominated by drought (Mittler, 2006; Rizhsky et al., 2002; Rizhsky et al., 2004; Sato et al., 2024; Zandalinas et al., 2016). Despite recognition of these interactions, quantitative responses of how legumes perform under combined heat and drought stress remains limited.

Actual-to-additive ratios have been used to establish the relationships between stresses. The method compares the observed trait value under combined heat and drought with the expected additive effect derived from single-stress treatments (Sadras et al., 2024; Sadras et al., 2025). Similar approaches have been applied in other crops and environmental contexts (Côté et al., 2016; Ploschuk et al., 2025), demonstrating the usefulness of additive frameworks for distinguishing antagonistic, additive, and synergistic responses (Sadras et al., 2024; Sadras et al., 2025; Ploschuk et al., 2025).

In this quantitative review, we integrated data from legume studies examining combined and individual drought and heat stress to evaluate developmental, physiological, and agronomic responses. Specifically, we (1) compiled published data from experiments assessing drought–heat relationships in legumes, (2) calculated actual-to-additive ratio to classify trait responses as additive, synergistic, or antagonistic, (3) identify sources of variability related to trait type, species, genotype, and experimental setting. From undertaking this approach we have been able to identify key research gaps that limit cross-species understanding and breeding applications, which could provide some direction for future research in addressing these gaps and challenges.

## 2. Methods

### 2.1 Literature review search

A literature search was conducted using the Web of Science database between the 31^st^ December 2024 and the 19^th^ February 2025, to identify studies investigating the combined effects of heat and drought stress on legumes. The following Boolean search string was used:

*“(Heat OR High temperature) AND (Drought OR Water Stress OR Water limited OR Water deficit) AND (Legume OR Soybean OR Glycine max OR Mungbean OR Vigna radiata OR Chickpea OR Cicer arietinum OR Medicago OR Lupin OR Faba OR Lentil OR Lens culinaris OR Peanut OR Arachis hypogaea OR Pea OR Lathyrus oleraceus OR Bean OR Cowpea OR Vigna OR Clover OR Serradella OR Alfalfa OR Lucerne OR Siratro OR Trifolium)”*.

To strengthen the search strategy, a manual review of reference lists and a search in additional databases were also undertaken. However, no additional studies were found beyond those retrieved from the initial database search.

The search was limited to research articles published in English, with no restriction on publication year. Titles and abstracts were screened for relevance, and only studies that applied factorial designs with at least all four treatment combinations of two water and two thermal regimes were retained. Studies that investigated heat or drought stress in isolation, or without a full factorial design, were excluded. Additionally, we retained pot-based studies in controlled environments, despite their recognised limitations, but excluded experiments where water deficit was induced with PEG (Cui et al., 2016; Passioura, 2006; Sadras et al., 2024).

### 2.2. Trait value extraction

Means and standard error estimates for plant traits were extracted from tables or digitised from figures using *WebPlotDigitizer* (v5.0). Extracted data included yield components, physiological traits (e.g. photosynthesis, transpiration) and phenological traits (e.g. time to flowering) and other traits reported by original studies (Table S1). Extracted traits were grouped into broad functional categories to facilitate analysis and interpretation. Carbohydrate metabolism included measurements of enzymes involved in starch and sugar metabolism, as well as metabolite levels. Oxidative stress and enzymatic defence encompassed traits reflecting antioxidant activity and the plant’s response to reactive oxygen species (ROS). Photosynthetic function included net photosynthetic rate, chlorophyll fluorescence, and other indicators of photosynthetic efficiency. Water/gas exchange captured traits related to the movement of water and gases through the plant, including processes that regulate transpiration, stomatal behaviour, and overall water status. Yield components comprised traits directly contributing to productivity, such as seed number, pod number, and biomass. Phenology included developmental timing traits, such as days to flowering or maturity. Traits that did not fit these categories, including morphological traits or nutrient content, were classified as other (Tables S1, S2).

### 2.3 Estimation of combined stress effects

To identify the relationship between heat and drought stress, we calculated actual-to-additive ratios, following the approach outlined by Sadras et al (2024, 2025). This approach tests the null hypothesis that when two stresses occur simultaneously, their combined effect is additive, i.e., it equals the sum of each stress acting independently. If the actual combined effect deviates from this total, it indicates that the stresses are interacting in a way that either enhances each other (synergistic) or one stress reduces the effect of the other (antagonistic).

The actual effects of heat stress (Δ*h*), water stress (Δ*w*) and combined heat and drought stress (Δ*c*) on plant trait (*T*) were calculated as differences from the control:

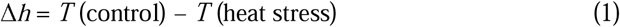

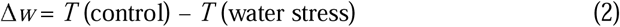

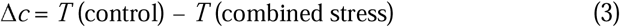

The null hypothesis of additivity (*A*) is:

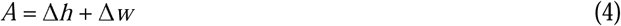

The actual-to-additive ratio is:

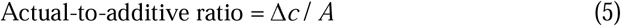

A ratio > 1 indicates a synergistic relationship, a ratio = 1 reflects an additive relationship, and a ratio < 1 suggests an antagonistic relationship. Applying the formulas described above, a total of 934 actual-to-additive ratios were calculated from the studies included in this review (Table 1).

**TABLE 1.**
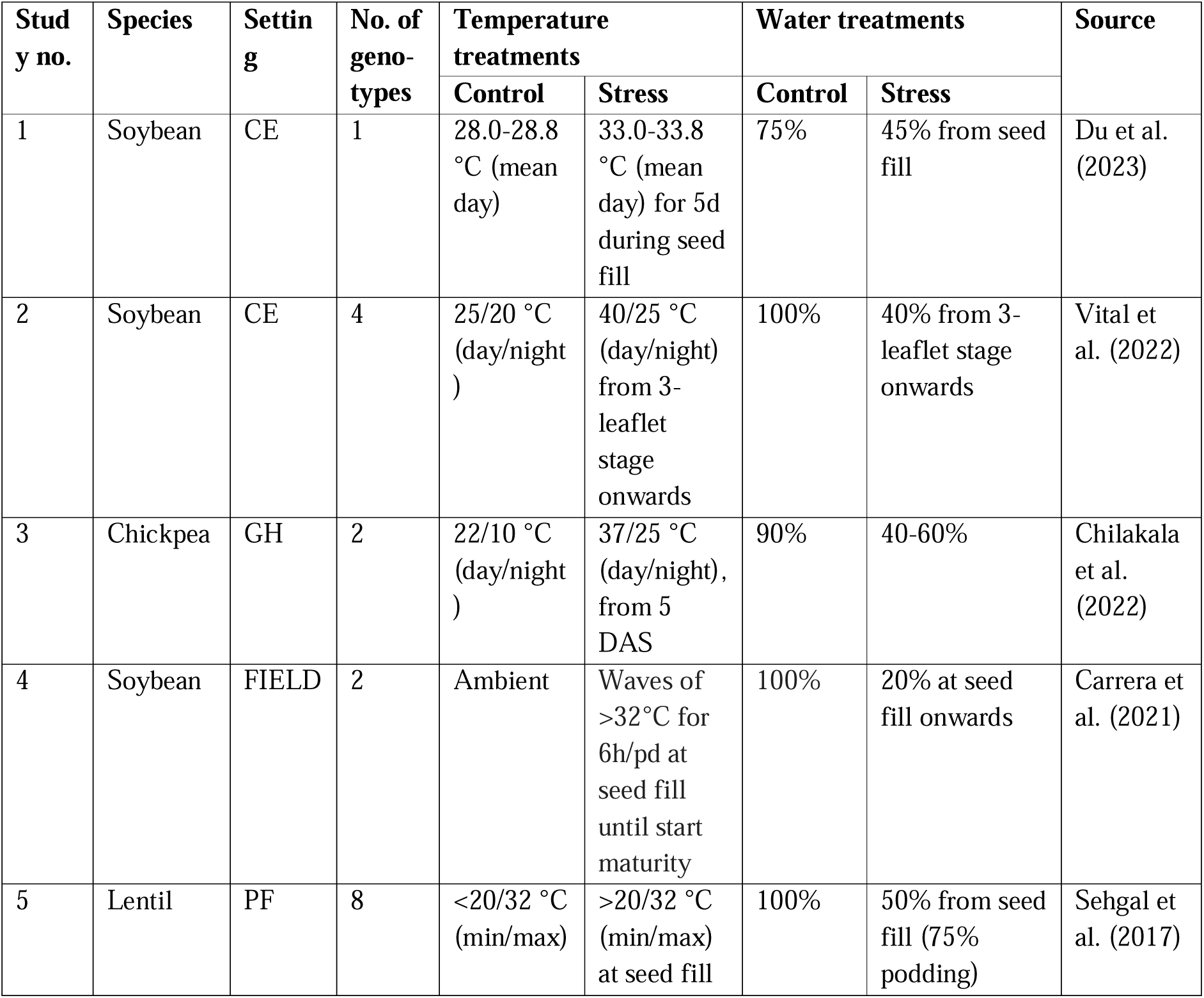
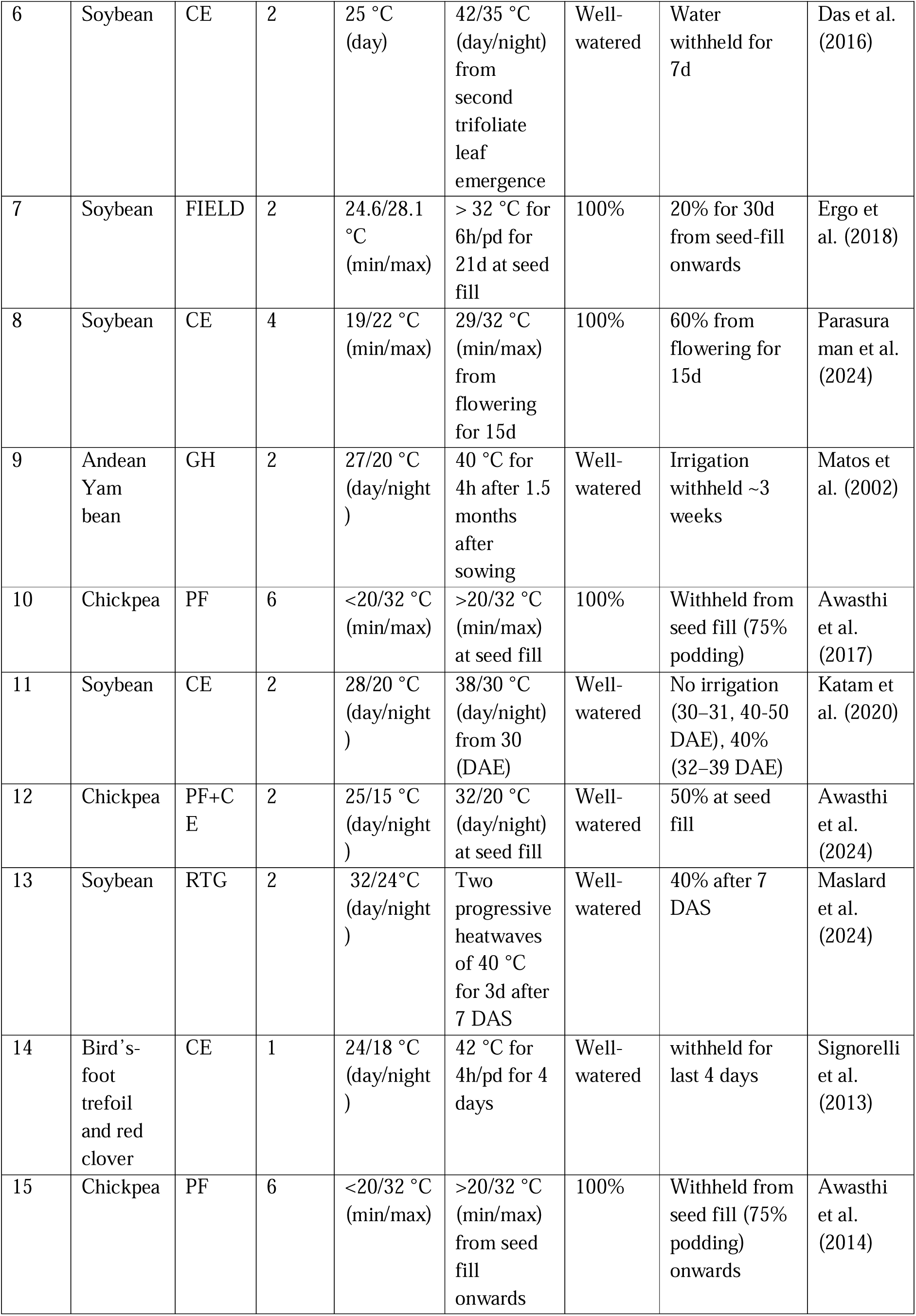
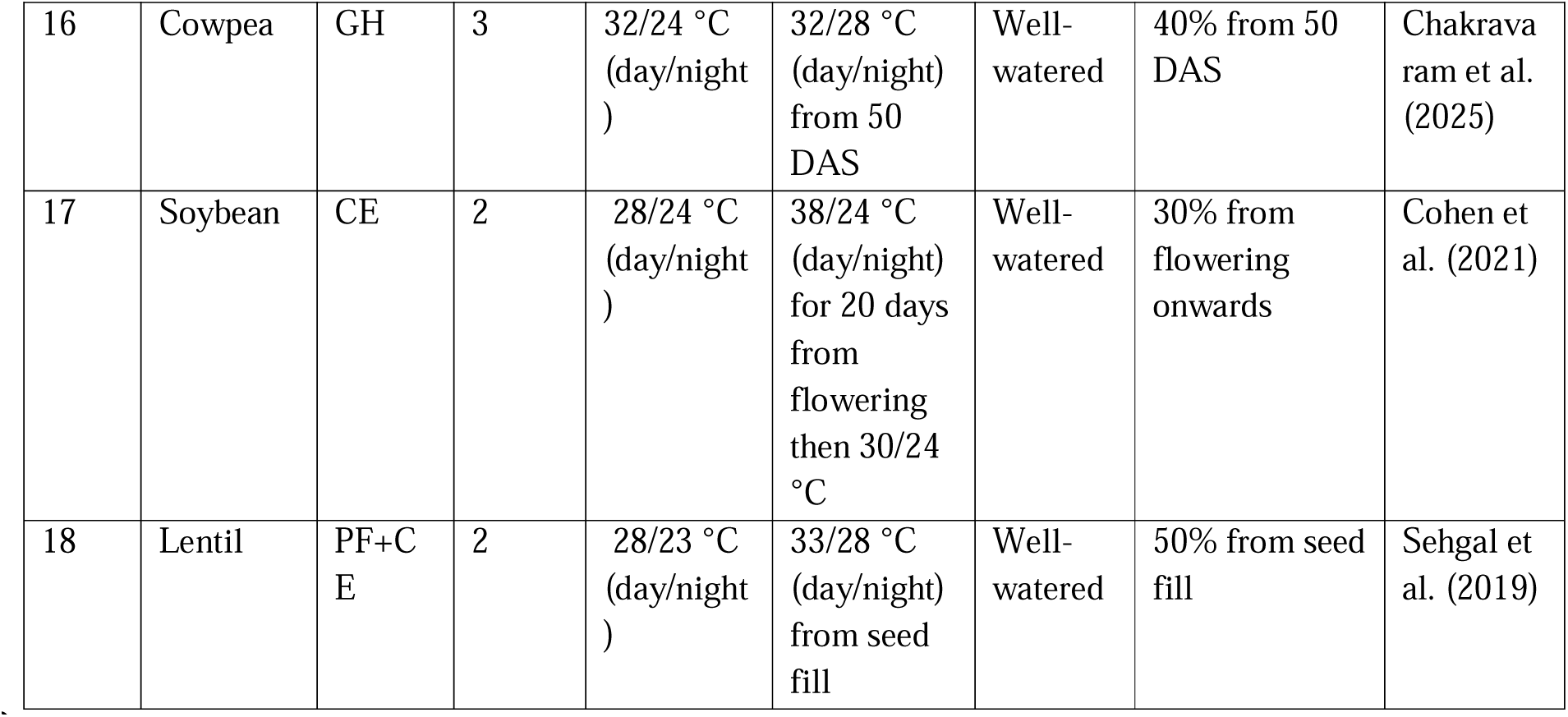
Summary of the 18 studies included in this review. The table presents the species investigated, experimental setting, number of genotypes assessed, and the temperature and water regimes applied under control and stress conditions. Experimental settings are potted plants in a greenhouse or glasshouse (GH), potted plants in controlled environment chambers (CE), potted plants in the field (PF), rhizotube in greenhouse (RTG), and field (FIELD). DAE refers to days after emergence and DAS corresponds to days after sowing. Percentages indicate soil water content as % of field capacity.

We analysed statistically the type of relationship against the null hypothesis of additivity with two complementary approaches: (1) the statistical analysis reported in the original paper including *p* for interactions from fitted statistical models and post hoc tests, and (2) actual-to-additive ratios. Where standard errors (SE) or standard deviations (SD) of means were reported, we propagated errors and calculated the 95% confidence interval CI for the actual-to-additive ratio (Table S3). This method assumes independent error terms; however, the individual effects of stresses were calculated using a common control. This implies that the calculated errors are not entirely independent, and consequently, the confidence intervals should be regarded as approximate. Furthermore, smaller means lead to large error of ratios because the standard error is inversely correlated to the mean (Sadras et al., 2024; Sadras et al., 2025). To avoid over-interpreting actual-to-additive ratios, we classified as ‘uncertain’ ratios that either lacked CI or with a CI greater than 1.0, as uncertainty of this magnitude exceeds the typical range of variation for additive, antagonistic, or synergistic interactions (Table S2). ATAR with CI ≤ 1.0 were classified according to their point estimate (< 1 antagonistic, = 1 additive, > 1 synergistic), and only actual-to-additive ratio values within the 0–2 range were considered to ensure biologically meaningful interpretation.

## 3. Results

### 3.1. A wide range of traits studied across species

A total of 18 studies investigating the combined effects of heat and drought stress on legumes were identified. These studies covered seven species, including chickpea (*Cicer arietinum*), lentil (*Lens culinaris*), soybean (*Glycine max*), cowpea (*Vigna unguiculata*), Andean yam bean (*Pachyrhizus ahipa*), bird’s-foot trefoil (*Lotus corniculatus*), and red clover (*Trifolium pratense*). A summary of the treatment conditions is provided in Table 1. The study number listed in the table is used as a reference for each study in the text, figures and tables.

Across all species, 929 trait responses were recorded, covering 130 distinct traits across the 18 studies. Most observations came from chickpea (332), lentil (274), and soybean (245), which together accounted for over 91% of all responses. Among all classified relationships, additive responses were the most common, accounting for 59%. Antagonistic responses were less frequent, representing 37% overall, while synergistic responses comprising only 4% of scores (Table 2).

**TABLE 2.** Number of responses to combined heat and drought stress classified as antagonistic, additive, synergistic, or uncertain, across seven legume species.

| Species | Antagonistic | Additive | Synergistic | Uncertain | Total |
| --- | --- | --- | --- | --- | --- |
| Chickpea | 110 | 105 | 10 | 107 | 332 |
| Lentil | 31 | 103 | 2 | 138 | 274 |
| Soybean | 25 | 44 | 3 | 173 | 245 |
| Cowpea | 10 | 16 | 0 | 25 | 51 |
| Andean yam bean | 3 | 4 | 10 | 7 | 14 |
| Bird's-foot trefoil | 2 | 2 | 0 | 3 | 7 |
| Red clover | 2 | 1 | 0 | 3 | 6 |
| **All species combined** | **19.7 %** | **29.6 %** | **1.6 %** | **49.1 %** | **929** |

For each trait category considered, responses were always mostly antagonistic (>40% of the responses), except for phenology (Table 3). Carbohydrate metabolism traits contributed the largest number of measurements (232), followed by oxidative stress and enzymatic defence-related traits (212), and photosynthetic function (185).

**Table 3.** Number of trait responses to combined heat and drought stress, classified as antagonistic, additive, synergistic or uncertain across trait categories.

| Trait category | Antagonistic | Additive | Synergistic | Uncertain | Total |
| --- | --- | --- | --- | --- | --- |
| Carbohydrate metabolism | 35 | 43 | 4 | 147 | 229 |
| Oxidative stress and enzymatic defence | 57 | 49 | 4 | 53 | 163 |
| Photosynthetic function | 26 | 56 | 1 | 101 | 184 |
| Water/gas exchange | 17 | 47 | 1 | 81 | 146 |
| Yield components | 33 | 28 | 2 | 42 | 105 |
| Phenology | 12 | 24 | 2 | 2 | 40 |
| Other | 3 | 28 | 1 | 30 | 61 |
| **Traits Combined** | **19.7 %** | **29.6 %** | **1.6 %** | **49.1 %** | **929** |

### 3.2 Yield

In study 1 (Du et al., 2023), a two-year pot experiment was conducted using the soybean cultivar Wandou 37. The plants were exposed to short-term heat of 5 days, long-term drought of 41 and 44 days in 2019 and 2020, respectively, and a combination of both conditions at seed fill. The results indicated that drought and the relationship between heat and drought had significant impacts on pod weight and seed weight per pod (*p* ≤ .05, based on least significant difference (LSD) tests in the original paper). In contrast, temperature exhibited weaker effects (*p* > 0.05; Du et al., 2021). Treatment effects became more evident between 19 and 33 days after treatment (DAT), when long-term drought and combined stress reduced both pod weight and seed weight per pod (Du et al., 2023). The actual-to-additive ratios indicate mild synergism at these time points for pod weight (1.18 at 19 DAT, 1.17 at 26 DAT, 1.20 at 33 DAT) and for seed weight per pod (1.10, 1.10, and 1.08, respectively), however, the associated CIs exceed the classification threshold, so these relationships cannot be assigned with certainty and were therefore considered uncertain (Table S2). These values, consistently above 1, suggest that heat stress amplified the drought-induced limitations on pod growth and seed filling. While drought was the dominant factor, its combination with heat intensified negative effects following stress exposure, leading to sustained reductions in pod weight (Du et al., 2023). Another soybean pot experiment, study 17 (Cohen et al., 2021), however, revealed antagonistic relationships for total seed weight and seed count (Fig. 1a). These contrasting outcomes may reflect differences in stress intensity and timing, as study 17 imposed heat stress approximately 5 °C higher than study 1 and during flowering rather than seed fill (Cohen et al., 2021; Du et al., 2023), suggesting that both the magnitude and developmental stage of heat exposure can shift the relationship between heat and drought on yield traits. A similar result was found in study 7 (Ergo et al., 2018), even though the soybean experiment was in a field setting and stress treatments varied (Fig. 1a).

**FIGURE 1.**
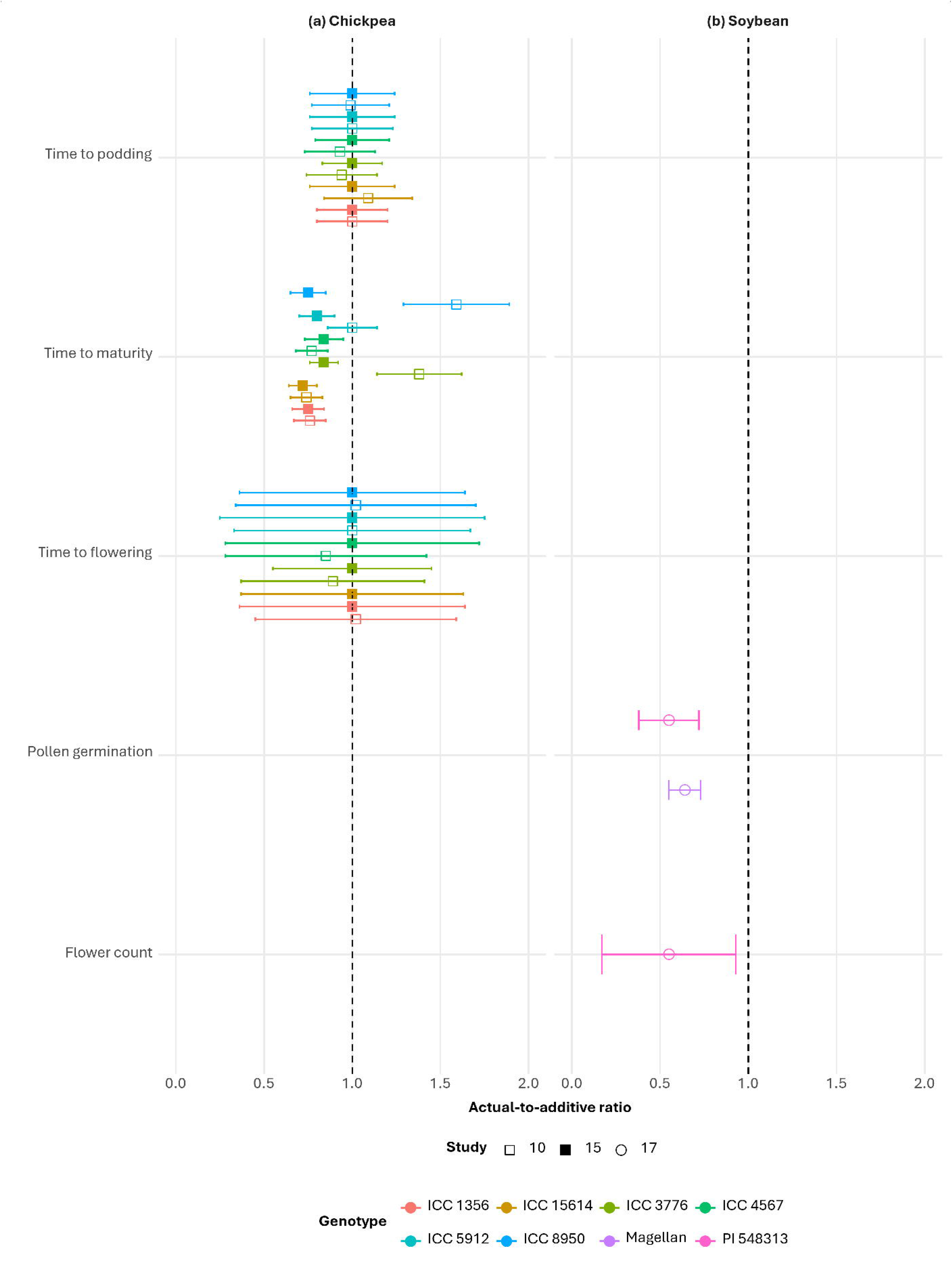
Actual-to-additive ratio for yield-related traits under combined heat and drought stress in (a) soybean, (b) chickpea and (c) lentil. Each point represents the actual-to-additive ratio (±95% confidence interval) for a given genotype and study, with colours denoting genotypes and symbols indicating study numbers. The dashed vertical line at 1.0 highlights the threshold for additive responses.

In studies 10 (Awasthi et al., 2017) and 15 (Awasthi et al., 2014), drought and heat were largely antagonistic for most chickpea yield traits, except for the filled pod-to-plant ratio, which was mostly additive, and genotypic differences were limited (Fig. 1b). Heat- or drought-tolerant genotypes (ICC 1356, ICC 15614, ICC 8950) produced more, heavier seed per plant when compared to their heat- or drought-sensitive counterparts (ICC 3776, ICC 4567, ICC 5912) in study 15 (*p* < 0.05; Awasthi et al., 2014), whereas the lack of genotypic differences in actual-to-additive ratio indicates the relationship between heat and drought stress was independent of tolerance.

In study 18 (Sehgal et al., 2019), lentil exhibited an additive response in total seed number and weight, individual seed weight and seed fill duration (Fig. 1c). In both study 1 (Du et al., 2023) with chamber-grown soybean and study 18 (Sehgal et al., 2019) with field-raised potted lentil plants that were moved to control chambers at time of stress, stress was applied during seed filling stage under comparable environmental conditions, with daytime temperatures around 33 °C and soil moisture maintained at 45–50% field capacity. Despite these similarities, the two species exhibited contrasting responses to combined heat and drought. This may reflect species-specific physiology or differences in experimental conditions, including pot size (55 × 60 cm vs. 20 × 15 cm), stress duration (5 vs. 10 days), and, where reported, relative humidity and radiation (both not reported in study 1; 60–65 % RH and 500 µmol m□² s□¹ in study 18).

### 3.3 Phenological traits

Few studies have examined phenological traits under combined heat and drought stress. Both field studies 10 (Awasthi et al., 2017) and 15 (Awasthi et al., 2014) applied stress at stage R5.5, which coincides with seed filling when 75 % of pods have formed. As a result, early phenological traits such as time to flowering or podding were not evaluated under heat and drought stress in chickpea, leading to additive relationships throughout (Fig. S1a). Time to maturity showed varying responses among genotypes. ICC 3776 and ICC 8950 exhibited antagonistic relationships in study 15, consistent with the other genotypes, but displayed synergistic responses under combined heat and drought stress in study 10 (Fig. S1a). Both studies were conducted at the same site by the same research group in different years, and slight differences in climatic conditions may explain these contrasting outcomes.

Study 17, which initiated stress at the onset of flowering, reported reductions in pollen germination and flower production in chamber-grown soybean (Cohen et al., 2021). However, these declines were less severe than expected based on the sum of individual stress effects, resulting in antagonistic relationships between heat and drought (Fig. S1b).

### 3.4 Leaf traits

In soybean, studies 1, 2, 6, 8, and 17 with potted plants in controlled growth chambers, and study 13 in a rhizotron, evaluated leaf traits such as chlorophyll fluorescence (Fv/Fm), leaf temperature, leaf water potential, net photosynthetic rate, stomatal conductance (Gs) and transpiration rate, across multiple cultivars (Fig. 2a). Across the six studies, which varied in experimental setting, heat and drought treatment intensity as well as in the duration and implementation of stress, there was substantial variation in the size and type of combined heat and drought effects on leaf traits. Of the 79 measurements across six traits and 15 genotypes, 52 (i.e. 66%) were classified as uncertain since they did not meet the requirements for classification outlined in the Methods. Out of the remaining 27 cases, additive relationships were most common, occurring 20 times (25%), while antagonistic and synergistic relationships were less frequent, with six (8%) and one (1%) instances, respectively.

**FIGURE 2.**
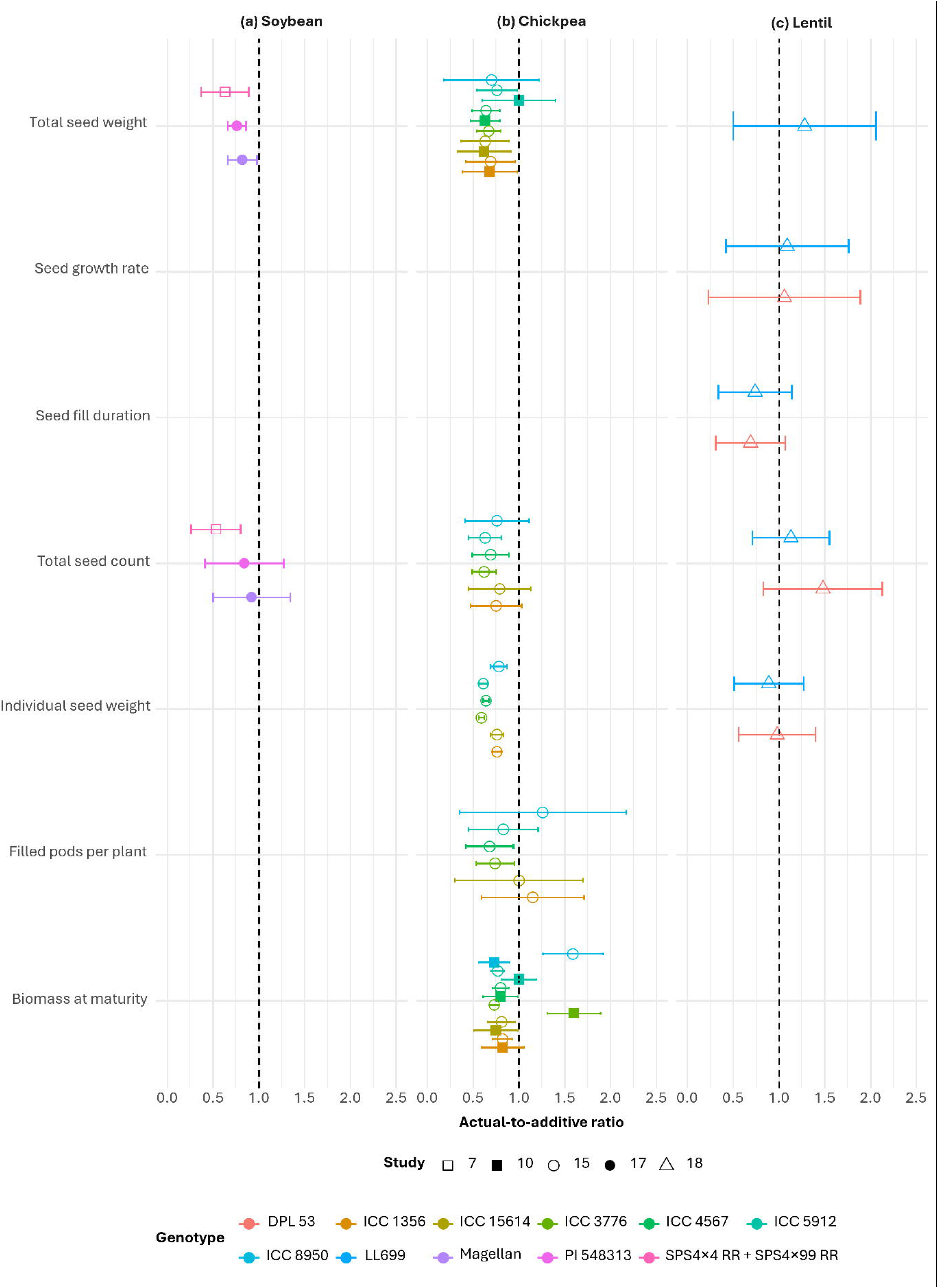
Actual-to-additive ratio for leaf traits under combined heat and drought stress in (a) soybean, (b) chickpea, (c) lentil, and (d) Andean yam bean. Each point represents the actual-to-additive ratio (±95% confidence interval) for a given genotype and study, with colours denoting genotypes and symbols indicating study numbers. The dashed vertical line at 1.0 highlights the threshold for additive responses.

In the most severe treatment applied to soybean, where the stress temperature reached 42 °C during the day and all water was withheld during the stress period of 7 days (Study 6; Das et al., 2016), synergy was observed for leaf water potential only in the genotype Surge (1.24±0.14), whereas the genotype Davidson exhibited antagonistic responses for this trait (0.76±0.06) as well as others. In a milder treatment with temperatures of 33.0-33.8 °C and irrigated to 45 % field capacity (Study 1; Du et al., 2023) produced an additive response for all classifiable traits. Several studies with intermediate heat stress (38–40 °C) and moderate soil moisture deficits (30–40 % field capacity; studies 2, 13, 17: Vital et al., 2022; Maslard et al., 2024; Cohen et al., 2021) showed mixed outcomes, with studies 2 and 13 producing strictly additive responses while study 17 also shows antagonistic relationships (Fig. 2a). Study 8 (Parasuraman et al., 2024), which applied a milder treatment (23–36 °C, 60 % field capacity), also recorded mixed responses, with eight synergistic and twelve antagonistic relationships (Table S2). However, because standard errors were not reported in the original study, confidence intervals could not be calculated, and these relationships are therefore classified as uncertain. These patterns suggest that greater stress intensity does not necessarily lead to stronger synergistic effects. Variation in actual-to-additive ratios was observed both between traits and between soybean genotypes within the same study. For instance, Pn and leaf temperature exhibited antagonistic responses in some studies, whereas Gs and Fv/Fm strictly showed additive outcomes (Fig. 2a). Together, these findings highlight that the magnitude and nature of combined stress effects are shaped by the interplay between trait, stress intensity, duration, timing and genotype.

In four studies (3, 10, 12, 15) focusing on chickpea (Fig. 2b), three of the previously mentioned leaf traits were evaluated under different experimental settings: study 3 was conducted in pots within a greenhouse (Chilakala et al., 2022), studies 10 and 15 used potted plants in the field (Awasthi et al., 2017; Awasthi et al., 2014), and study 12 involved potted plants grown in the field that were later transferred to controlled chambers at seed fill to impose stress (Awasthi et al., 2024). Among 21 scores, antagonistic relationships were the most prevalent, occurring 11 (52%) times, followed by additive relationships (nine occurrences, i.e. 43%) and synergistic relationships with only one occurrence (5%; Fig. 2b). Of the 8 genotypes examined, six (ICC 1356; heat-tolerant, ICC 15614; heat-tolerant, ICC 3776; drought-sensitive, ICC 5912; heat-sensitive, ICC 8950; drought-tolerant, and ICC 4567; heat-sensitive) were included in both studies 10 (Awasthi et al., 2017) and 15 (Awasthi et al., 2014). The environmental conditions in both pot-in-field studies were similar, featuring early and late sowing to induce heat stress, while drought stress was introduced by withholding water at seed filling when plants reached 75% podding. Caution is needed in the comparison because experiments were conducted in two different seasons, where seasonal factors such as radiation, vapour pressure deficit and wind might have altered the relationships. Furthermore, ICC 3776 and ICC 8950 were also part of study 12 (Awasthi et al., 2024), which followed a similar experimental setup; however, in this instance, the stressed pots were moved into controlled chambers rather than remaining outside.

Heat-tolerant chickpea genotypes such as ICC 1356 and ICC 15614 exhibited additive responses to Fv/Fm, but were antagonistic for Gs across studies. Heat-sensitive genotypes, including ICC 4567 and ICC 5912, responded almost entirely with antagonistic relationships for either trait (Fig. 2b). In contrast, study 15 (Awasthi et al., 2014) found that tolerant genotypes maintained greater Gs and improved Fv/Fm when exposed to combined heat and drought stress, in comparison to sensitive genotypes (LSD test, *p* < 0.05). For genotypes ICC 3776 and ICC 8950, antagonistic outcomes were exclusively observed for both traits in study 15, while study 10 recorded a synergistic result for ICC 3776 (1.99±0.7) and an additive response for ICC 8950 in Fv/Fm (1.11±0.27), and study 12 found that ICC 3776 responded additively (1.27±0.99) for Fv/Fm. These contrasting outcomes illustrate that genotype responses can differ even under similar environmental conditions.

Study 3 investigated the impact of combined heat and drought stress on the occurrence and severity of dry root rot disease caused by *Macrophomina phaseolina* in chickpea, using potted plants grown in a greenhouse. The combined stresses reduced both root and leaf water potential to a greater extent than each individual stressor for the two genotypes tested. In many instances, both traits also exhibited a significant reduction when subjected to combined stress compared to the effects of a single stressor in conjunction with pathogen treatment (*p* < 0.05; Chilakala et al., 2022). However, the drought-tolerant genotype ICC 4958 and JG 62, which is susceptible to dry root rot and sensitive to salinity, both showed additive responses for leaf and root water potential (Fig. 2b; Table S1).

In lentil (Fig. 2c), Fv/Fm, Gs, and Pn were measured in two studies under comparable conditions (Sehgal et al., 2019; Sehgal et al., 2017). Study 5 assessed eight genotypes, while study 18 included two, both sharing the drought-tolerant DPL 53. All genotypes responded additively (where classifiable) to Fv/Fm, Gs and Pn in both studies 5 and 18 (Fig. 2c). Study 9 (Matos et al., 2002) measured Fv/Fm, Pn and Gs in Andean yam bean using two genotypes. After applying the CI threshold, the Fv/Fm value for genotype AC 102 and both Gs measurements could not be classified (Table S2). Genotype AC 524 showed an antagonistic Fv/Fm response. Pn responses were additive across genotypes (Fig. 2d).

Other traits related to photosynthesis and gas exchange, including chlorophyll concentration and relative water content, exhibited a comparable pattern in response to heat and drought stress: relationships were additive in 55 (71%) out of 77 classified cases, antagonistic in 22 (29%) and none were synergistic (Table S1).

### 3.5 Oxidative stress and enzymatic defence

Heat and drought trigger pronounced oxidative stress in legumes, evident from elevated reactive oxygen species (ROS) and downstream markers of cellular damage (Hussain et al., 2019; Wang et al., 2024). In chickpea study 10 (Awasthi et al., 2017), hydrogen peroxide (H□O□) increased under combined stress, though the magnitude and relationship varied by genotype and tissue. Overall, leaf tissue accumulated more H□O□ than seeds, with stress-sensitive genotypes exhibiting higher levels compared to tolerant ones (Awasthi et al., 2017). Heat-tolerant and heat-sensitive genotypes ICC 1356, ICC 4567 and ICC 5912 demonstrated an additive accumulation of H□O□ in chickpea leaves and seeds, while heat-tolerant genotype ICC 15614 experienced synergistic responses (Fig. 3a). In contrast, all drought-tolerant and drought-sensitive genotypes primarily displayed an antagonistic response across tissues (Fig. 3a).

**FIGURE 3.**
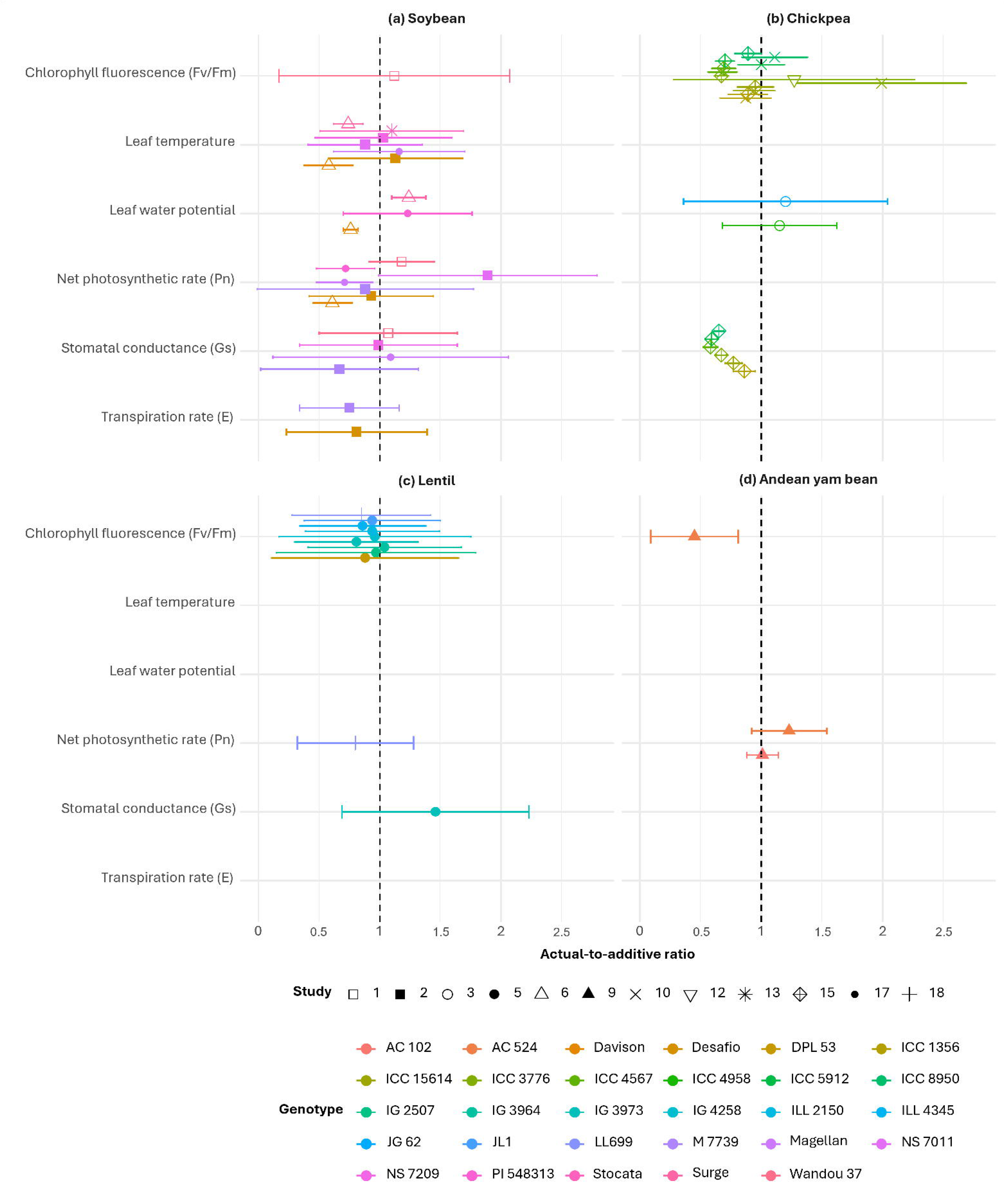
Actual-to-additive ratio for stress indicators under combined heat and drought stress in (a) chickpea and (b) lentil. Each point represents the actual-to-additive ratio (±95 % confidence interval) for a given genotype and study, with colours denoting genotypes and symbols indicating study numbers. The dashed vertical line at 1.0 highlights the threshold for additive responses.

Potential oxidative damage induced by reactive oxygen species (ROS) can be quantified through malondialdehyde (MDA) accumulation, which reflects lipid peroxidation, and by measuring electrolyte leakage as an indicator for programmed cell death (Demidchik et al., 2014; Jambunathan, 2010; Khaleghi et al., 2019; Zhang et al., 2021). Across legume species, oxidative stress responses to combined heat and drought were highly variable, with both tissue type and genotype strongly influencing outcomes.

In chickpea (study 10), MDA was consistently higher in leaves than in seeds across all genotypes, and although genotype, treatment, and their interaction were significant (Awasthi et al., 2017), yet actual-to-additive ratios revealed that most responses were additive or mildly antagonistic (Fig. 3a). Electrolyte leakage in chickpea across studies 10, 12 and 15 produced mainly additive outcomes under combined stress, except two genotypes in study 15 that yielded antagonistic outcomes (Fig. 3a; Awasthi et al., 2024; Awasthi et al., 2017; Awasthi et al., 2014). Environmental variation between trials in studies 10 and 15 contributed to inter-genotypic differences in electrolyte leakage, highlighting the influence of subtle weather factors on stress responses.

In lentil, studies 5 (Sehgal et al., 2017) and 18 (Sehgal et al., 2019), both conducted by the same research group under differing environmental conditions (Fig. 3b), assessed multiple genotypes for electrolyte leakage. All five measurements were found to be additive. Results with higher actual-to-additive ratios were found in study 5 where pots remained in the field throughout the trial. However, due to large CI, these ratios could not be accurately classified.

To mitigate oxidative stress, plants activate enzymatic antioxidants (superoxide dismutase, SOD; glutathione reductase, GR; ascorbate peroxidase, APX; catalase, CAT and peroxidase, POX), to actively detoxify ROS (Gill & Tuteja, 2010; Hasanuzzaman et al., 2020). Across four studies encompassing 14 genotypes, 53 classifiable measurements were reported for these traits. Antagonistic relationships were predominant, occurring in 34 cases (i.e. 64%), followed by additive (17, 32%) and synergistic (2, 4%), indicating that combined stress rarely amplifies antioxidant activity beyond what is expected from individual stresses (Fig. 4).

**FIGURE 4.**
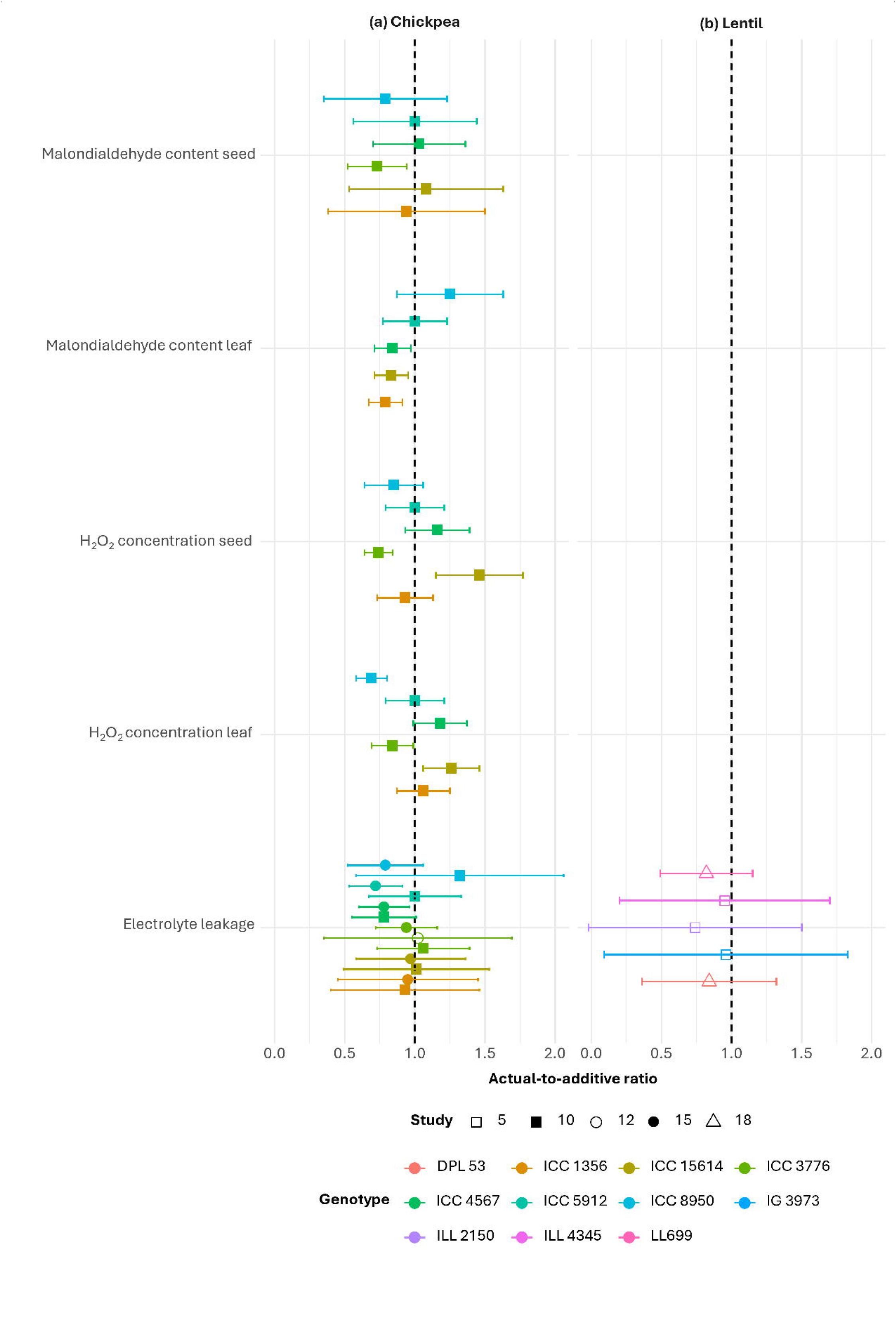
Actual-to-additive ratio for enzymatic oxidants under combined heat and drought stress in (a) chickpea, (b) clover, (c) bird’s-foot trefoil and (d) soybean. Each point represents the actual-to-additive ratio (±95% confidence interval) for a given genotype and study, with colours denoting genotypes and symbols indicating study numbers. The dashed vertical line at 1.0 highlights the threshold for additive responses.

Genotypic differences were evident, particularly in chickpea study 10 (Awasthi et al., 2017) and soybean study 11 (Katam et al., 2020), as shown in Figure 4a,d. Tissue type, however, appeared to have minimal influence on enzyme activity in chickpea (Fig. 4a). Clover and bird’s-foot trefoil (Study 14) demonstrated similar relationships with SOD under combined stress (Fig. 4b,c; Signorelli et al., 2013).

### 3.6 Carbohydrates

Carbohydrates, and their associated metabolic enzymes, coordinate source–sink carbon allocation and often increase during abiotic stress that decouple tissue expansion and photosynthesis (Muller et al., 2011). Across five studies, lentil (studies 5 and 18; Sehgal et al., 2019; Sehgal et al., 2017) and chickpea (studies 12 and 15; Awasthi et al., 2024; Awasthi et al., 2014) showed mainly additive and antagonistic responses (Fig. 5a,b). Notably, across the species, reducing sugar concentration in leaf and seed tempted to respond antagonistically while the other traits saw more variation. Soybean results (study 1; Du et al., 2023) were limited to an additive response (1.03±0.33) in starch content in the leaves (Fig. 5c).

**FIGURE 5.**
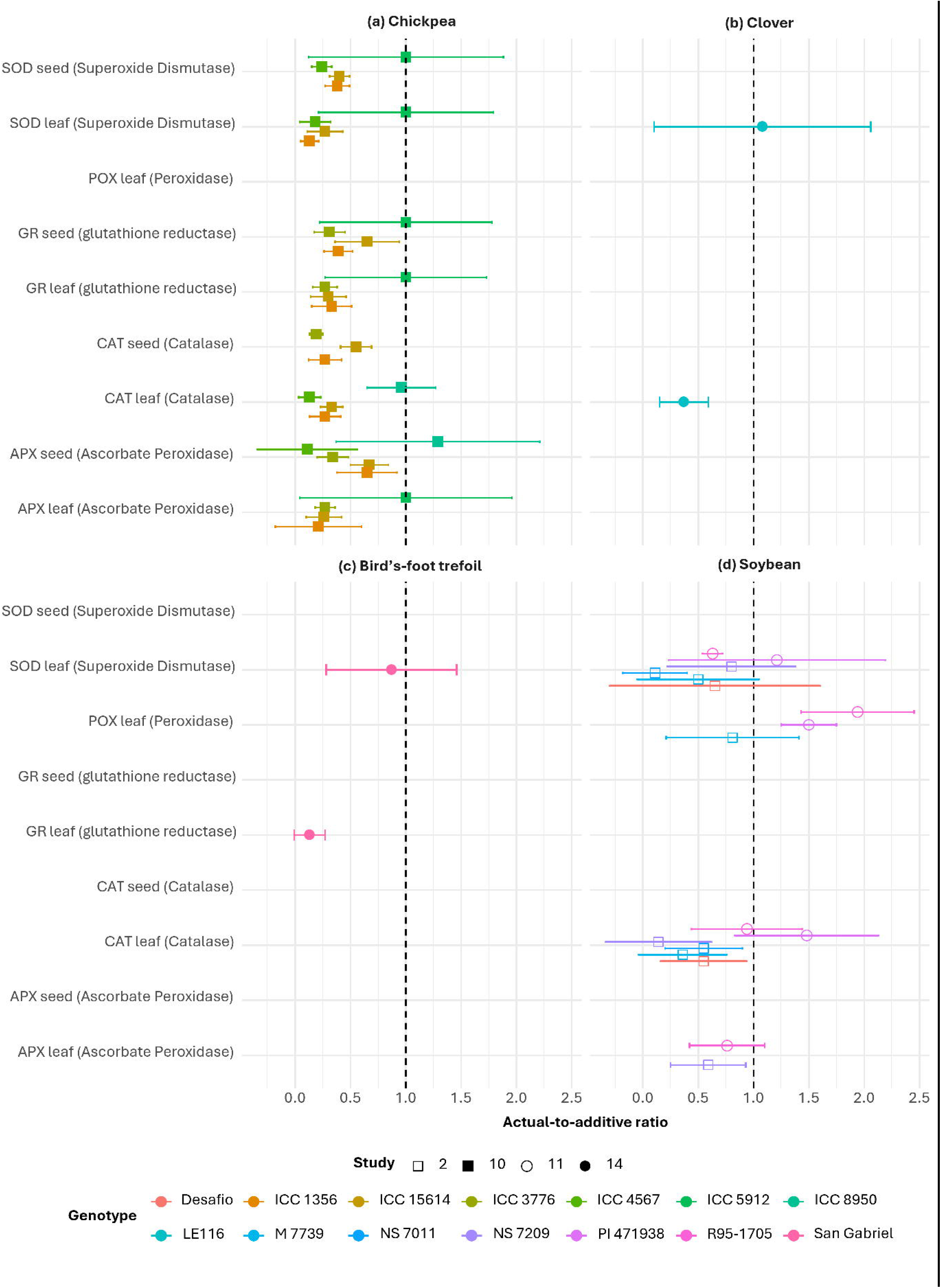
Actual-to-additive ratio for carbohydrates and metabolic enzymes under combined heat and drought stress in (a) lentil, (b) chickpea and (c) soybean. Each point represents the actual-to-additive ratio (±95% confidence interval) for a given genotype and study, with colours denoting genotypes and symbols indicating study numbers. The dashed vertical line at 1.0 highlights the threshold for additive responses.

### 3.7 Night-time heat stress in cowpea

In study 16, Chakravaram et al. (2025) examined three cowpea genotypes under combined night-time heat and drought stress imposed from flowering onwards. Night temperature was increased by four degrees, resulting in 32 °C day and 28 °C night conditions, while irrigation was reduced to 40% of the control. The genotypes EpicSelect.4, LOUVI (reported as LOVI by Chakravaram et al., 2025) and UCR 369 were chosen for their similar phenology but contrasting waterlogging responses as described by Olorunwa et al. (2022). Across 13 traits and 26 classified traits, 16 relationships were additive (62%), ten antagonistic (38%) and none synergistic. Antagonistic effects were most evident in physiological traits related to pigment content and photosystem efficiency, while yield traits exclusively displayed additive responses. Yield-related traits exhibited greater genotypic and treatment variation than other traits in the original study (*p* < 0.05; Chakravaram et al., 2025), suggesting that night-time stress amplified the impact of drought on yield more than on physiological function. This aligns with the higher actual-to-additive ratios in yield-related traits; however, differences in ratios among genotypes appear to be limited (Fig. 6).

**FIGURE 6.**
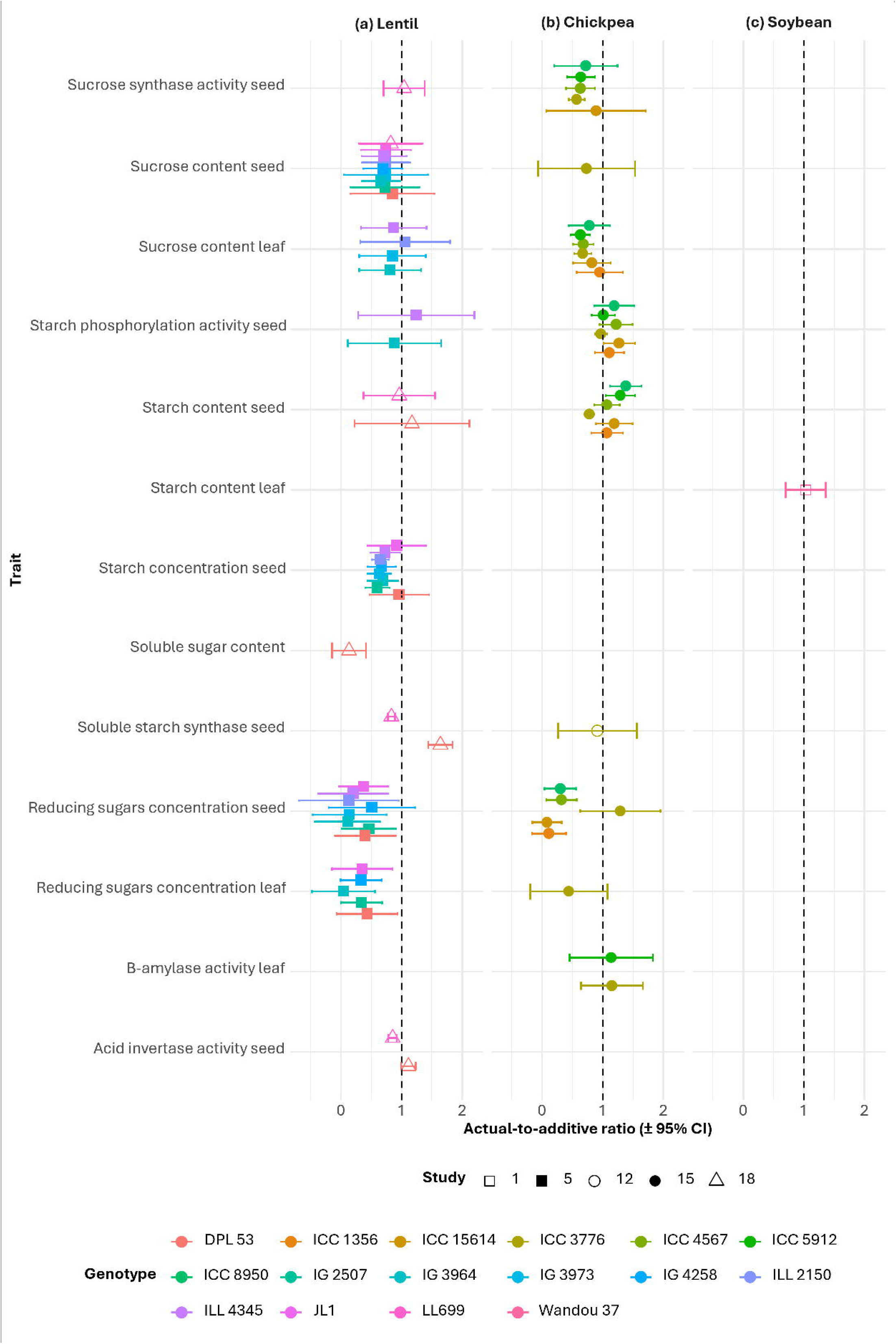
Actual-to-additive ratio for plant traits under combined nighttime heat and drought stress in cowpea. Each point represents the actual-to-additive ratio (±95 % confidence interval) for a given genotype, with colours denoting genotypes. The dashed vertical line at 1.0 highlights the threshold for additive responses.

## 4. Discussion

### 4.1 Patterns of combined stress response

Understanding how legumes respond to concurrent heat and drought stress is critical for crop yield under climate change. Our analysis demonstrates that combined stress were mostly antagonistic or additive, with the prevalence and direction of responses influenced by species, genotype, trait, and environmental and experimental context. Approximately 49% of the 929 trait responses were classified as uncertain, reflecting insufficient precision to determine whether combined heat and drought effects were antagonistic, additive, or synergistic. This substantial proportion indicates that many current experimental approaches lack the resolution needed to reliably characterise stress relationships (e.g. limited replication and inconsistent stress imposition). Within the remaining classifiable responses, clear patterns emerged: additive relationships were most common, accounting for 59% of observations, followed by antagonistic responses at 37% and synergistic effects at 4% (Tables 2a,b and S1). Evidence from other systems supports this pattern. In Ploschuk et al. (2025), examining waterlogging and heat stress, synergy was observed in only 4 to 9% of traits. Similarly, out of 329 relationships between drought and other factors (e.g. nutrients, CO_2_ concentration) affecting plant and aphid traits, 55% were additive, 36% antagonistic, and 9% synergistic (Sadras et al., 2025). No synergies were reported for virus–drought relationships affecting plant traits (Sadras et al., 2024). These observations are consistent with the conclusion that synergies are rare, context-specific, and do not scale from lower (molecular) to higher (population, ecosystem) levels, whereas additive and antagonistic relationships dominate under multiple stress conditions, challenging the common assumption that concurrent stresses inevitably exacerbate damage (Côté et al., 2016).

### 4.2 Limitations of experimental approaches

Several limitations constrain the reliability of experimental studies on combined stresses. Most published studies relied on potted plants grown in controlled environments such as glasshouses or chambers, which restrict root development, modify canopy architecture, and alter light and temperature regimes relative to field conditions (Annunziata et al., 2017; Aphalo & Sadras, 2021; Freschet et al., 2021; Passioura, 2006; Sellaro et al., 2024). The imposition of stress frequently occurred abruptly or intensely, which restricts plant adaptation and might have amplified or distorted the effects of the imposed abiotic stressors (Das et al., 2016; Passioura, 2006; Signorelli et al., 2013). For example, in soybean, study 2 (conducted in Brazil) imposed heat stress at 40 °C for eight days, while study 6 (from Canada) maintained plants at 42 °C for seven days, both in controlled growth chambers (Das et al., 2016; Vital et al., 2022). Such sustained and extreme temperatures are unlikely to represent realistic field conditions, particularly when applied continuously in pots.

Building on these limitations, gaps persist that constrain our understanding of legume responses to combined heat and drought stress. Research remains heavily skewed toward a few species, primarily grain legumes such as chickpea, lentil, and soybean, whereas other grain legumes (e.g. mungbean, cowpea, faba bean, common bean, Andean yam bean) and pasture legumes (e.g. lucerne, white clover, red clover, birds-foot trefoil) remain largely understudied (Table 2). Genotype-specific responses were common, yet the diversity of genotypes tested was uneven across species, and differences in experimental setting and environment likely contributed to variation between experiments.

### 4.3 Underexplored roles of high night temperature stress

The impacts of elevated night temperature on yield have been documented in several crops including wheat (García et al., 2018; Hein et al., 2020; Prasad et al., 2008), barley (García et al., 2018), rice (Kumar et al., 2021), maize (Hein et al., 2024), quinoa (Lesjak & Calderini, 2017), and cotton (Loka & Oosterhuis, 2010), yet few studies have explored its combined effect with drought. The results of Chakravaram et al. (2025) indicate that the nature of these relationships is genotype- and trait-specific, highlighting the importance of breeding for adaptation to both daytime and night-time thermal stress co-occurring with drought.

### 4.4 Trait coverage and stress quantification

Our study covered 130 unique traits, yet relatively few were measured consistently across multiple species or studies (Table S1). Temporal and developmental dimensions of stress remain poorly explored, as few studies impose stress across multiple growth stages or directly compare early vegetative versus reproductive responses, limiting insight into stage-specific relationships. This gap is particularly significant for legumes, many of which exhibit indeterminate growth, with flowering, pod filling, and vegetative development overlapping in time (Ambika et al., 2021; Dudley et al., 2025). Consequently, the timing, duration, and recurrence of stress events can differentially affect organs at distinct developmental stages, complicating both physiological interpretation and breeding for adaptation to combined stresses (Devi et al., 2023; Wang et al., 2006). Few studies in our dataset assessed phenological traits because stress treatments were typically imposed at fixed growth stages; however, incorporating such traits could offer valuable insight into adaptive responses to combined stress.

In addition, inconsistencies in quantifying stress intensity and duration hinder cross-study comparisons and the development of predictive models. In many cases, stress treatments are nominal, with no quantification of plant stress, e.g. measuring leaf expansion rate or leaf water potential. Together, these limitations show that even though many traits have been measured, our understanding of how legumes respond to combined heat and drought remains incomplete and heavily dependent on experimental conditions.

Advancing legume production under concurrent heat and drought stress requires multi-species, multi-genotype evaluations across different developmental stages, combined with factorial manipulation of stress intensity and timing under standardised, robust protocols. Field-based studies under realistic fluctuations of temperature and radiation are crucial for agronomic relevance. Owing to niche-construction, namely the effect of organisms in their own niche, the niche of others or both (Odling-Smee, 2024; Sadras & Hayman, 2025), agronomically relevant traits including yield and photosynthesis rarely scale from single plant (Pettigrew et al., 1989). Reliable protocols for field experiments (Sinclair, 2022) will support breeding strategies that focus on key trait combinations, such as stress tolerance at critical growth stages and maintenance of yield stability, helping to identify legume cultivars that are genuinely adapted to the simultaneous challenges of heat and drought. Coupling these experimental frameworks with modelling approaches will be critical for extrapolating insights beyond measured conditions, predicting crop performance under future climates, and ultimately guiding breeding and management decisions in multi-stress environments.

## Author Contributions

Lotus Meijer: conceptualisation (equal); methodology (supporting); data curation (lead); formal analysis (lead); writing – original draft (lead); writing – review & editing (equal). Shanice Van Haeften: conceptualisation (equal); methodology (supporting); data curation (supporting); supervision (lead); writing – review & editing (equal). Karine Chenu: conceptualisation (equal); methodology (supporting); writing – review & editing (equal). Millicent R. Smith: conceptualisation (equal); methodology (supporting); funding acquisition (lead); writing – review & editing (equal). Victor Sadras: conceptualisation (equal); methodology (lead); Writing – review & editing (equal).

## Acknowledgements

This work was partially funded by Grains Research and Development Corporation (GRDC) as part of project UOQ2402-010RTX ‘Fast tracking deployment of chickpea heat tolerance to develop chickpea varieties with improved high temperature tolerance’. We acknowledge support from the Deutsche Forschungsgemeinschaft (DFG) and University of Queensland International Research Training Group 2843 – Accelerating Crop Genetic Gain.

## Data Availability Statement

All data were sourced from published literature.

---

**FIGURE S1.**
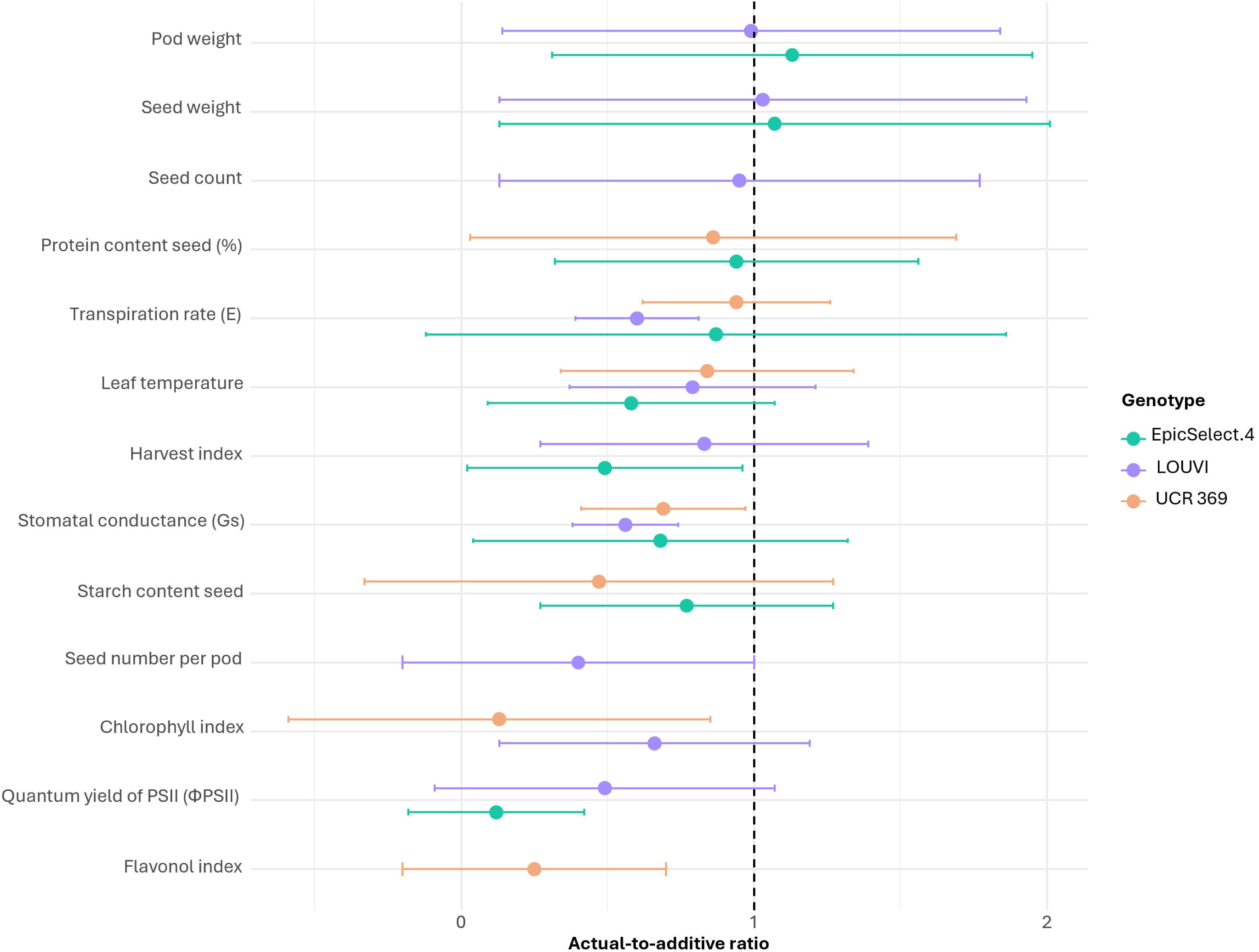
Actual-to-additive ratio for phenology traits under combined heat and drought stress in (a) chickpea and (b) soybean. Each point represents the actual-to-additive ratio value (±95 % confidence interval) for a given genotype and study, with colours denoting genotypes and symbols indicating study numbers. The dashed vertical line at 1.0 highlights additive responses.

